# *Memo1* gene expression in kidney and bone is unaffected by dietary mineral load and calciotropic hormones

**DOI:** 10.1101/2020.02.06.933960

**Authors:** Matthias B. Moor, Olivier Bonny

**Author notes:** Correspondence to: Matthias B. Moor, MD PhD, Department of Nephrology and Hypertension, Bern University Hospital, Freiburgstrasse 15, 3010 Bern, Switzerland.

## Abstract

Mediator of Cell Motility 1 (MEMO1) is an ubiquitously expressed modulator of cellular responses to growth factors including FGF23 signaling, and *Memo1*-deficient mice share some phenotypic traits with *Fgf23*- or *Klotho*-deficient mouse models. Here, we tested whether *Memo1* gene expression is regulated by calciotropic hormones or by changing the dietary mineral load.

MLO-Y4 osteocyte-like cells were cultured and treated with 1,25(OH)_2_-vitamin D_3_. Wildtype C57BL/6N mice underwent treatments with 1,25(OH)_2_-vitamin D_3_, parathyroid hormone (PTH), 17β-estradiol or vehicle. Other cohorts of C57BL/6N mice were fed diets varying in calcium or phosphate content. Expression of *Memo1* and control genes was assessed by qPCR.

1,25(OH)_2_-vitamin D_3_ caused an acute decrease in *Memo1* transcript levels in vitro, but not in vivo. None of the hormones tested had an influence on *Memo1* transcripts, whereas the assessed control genes reacted the expected way. Dietary interventions with calcium and phosphate did not affect *Memo1* transcripts but altered the chosen control genes’ expression.

We observed that *Memo1* was not regulated by calciotropic hormones or change in mineral load, suggesting major differences between the regulation and physiological roles of *Klotho, Fgf23* and *Memo1*.

## Introduction

*Memo1* is an evolutionary conserved protein in all kingdoms of life that has shown intracellular expression in cytoplasma and nucleus (Schlatter et al., 2012, Haenzi et al., 2014, Moor MB, 2018). A conditional and inducible knockout mouse model with postnatal deletion of exon 2 of the *Memo1* gene has resulted in a syndrome of premature aging with elevated calcemia, elevated FGF23, insulin hypersensitivity and bone disease (Haenzi et al., 2014, Moor, 2018). These traits share a partial overlap with mouse models deficient in KLOTHO (Kuro-o et al., 1997) or FGF23 (Shimada et al., 2004), two proteins which have tremendously reshaped our understanding of the regulation of calcium and phosphate metabolism by the intestine, kidney and bone (Hu et al., 2013, Moor and Bonny, 2016).

In addition, evidence from cell culture experiments investigating co-localization and phosphorylation status of adaptor proteins suggested that MEMO1 protein participates in and modulates a signaling cascade involving FGF23 and the FGFR (Haenzi et al., 2014).

Serum analyses of *Klotho* and *Fgf23*-deficient mice showed excessive 1,25(OH)_2_-vitamin D_3_ levels, a finding that has been variably found in *Memo1*-deficient mice depending on the genetic background (Haenzi et al., 2014, Moor, 2018). The promoters of *Fgf23* and *Klotho* both contain vitamin D response elements (Forster et al., 2011, Orfanidou et al., 2012). FGF23 secretion is increased by parathyroid hormone (PTH) (Lavi-Moshayoff et al., 2010) and 17β-estradiol (Carrillo-Lopez et al., 2009). Regarding the regulation of *Memo1*, a transcriptomic analysis of rat pineal glands has detected increased *Memo1* transcripts upon synthetic estrogen treatment compared to controls (Deffenbacher, 2006), highlighting a potential endocrine regulation of the transcription of the *Memo1* gene. Here, we therefore tested the hypothesis that *Memo1* expression can be regulated by minerals or calcitropic stimuli.

## Methods

### Cell culture

The mouse long bone osteocyte-Y4 cell line (MLO-Y4) was kindly provided by Lynda Bonewald (Kato et al., 1997). MLO-Y4 cells were maintained in culture in minimal essential medium alpha-modified, alpha-MEM (Gibco by Life Technologies, Carlsbad, CA, USA), containing 2.5% heat-inactivated calf serum (Sigma), 2.5% heat-inactivated fetal bovine serum (Sigma), and 1% penicillin/streptomycin (Invitrogen by Life Technologies). Serum heat inactivation was carried out in water bath at 56°C for 30min. Cells were cultured on rat-tail type I collagen (Invitrogen by Life Technologies). For vitamin D stimulation, cells were kept in serum-free medium supplemented with 10nM 1,25(OH)_2_-vitamin D_3_ (Sigma) or ethanol vehicle for 24h.

### Animal experiments

C57BL/6N mice were obtained from Janvier (Le Genest-Saint-Isle, France). Mice were held in a conventional animal facility with up to 6 animals per cage, and they were fed a standard laboratory chow (Kliba Nafnag TS3242; calcium 1%, phosphorus 0.65%, magnesium 0.23%, vitamin D 1600 IU/kg, vitamin A 27000 IU/kg, vitamin E 150mg/kg, protein 18.8%, crude fat 5.6%, crude fibre 3.5%, lysine 1.1%; KLIBA Kaiseraugst, Switzerland) unless specified otherwise and were kept on 12/12 (experimental) or 14/10 (breeding) light-dark cycles. All animal experimental protocols were approved by the State Veterinary Service of the Canton de Vaud, Switzerland. For all mouse studies, sample sizes were considered based on previous results in our laboratory.

### Dietary interventions

For studies of mineral metabolism, specifically designed diets and hormones were used. Calcium diet challenge experiments were carried with 5 male C57BL/6N mice per condition in their home cage, all aged 12 weeks. Mice were randomly allocated to be fed modifications of KLIBA 2222 diet containing either 0.17% (low calcium diet), 0.82% (normal calcium diet) or 1.69% (high calcium diet) (KLIBA, 2222) over 7 days. All calcium diets contained phosphorus 0.8%, vitamin A 4000 IU/kg, vitamin D 1000 IU/kg, vitamin E100 mg/kg, protein 18%, crude fat 7%, lysine 14g/kg (Kliba, 2222). For a phosphate diet challenge, 3 groups of 5 male C57BL/6N mice all aged 13-14 weeks were kept in metabolic cages and randomly allocated to be fed diets containing low (0.2%), intermediate (0.8%), or high (1.5%) phosphate content over 7 days. The phosphate diets were modifications of KLIBA 2222 diet (calcium 1.2%, vitamin A 4000 IU/kg, vitamin D 1000 IU/kg, vitamin E100 mg/kg, protein 18%, crude fat 7%, lysine 14g/kg).

### Hormone injections

All hormone injections were performed in the home cage of the mice after random treatment allocation of individual ear-marked mice. For 1,25(OH)_2_-vitamin D_3_ treatment, male C57BL/6N mice aged 13-15 weeks were subcutaneously injected with 2 μg/kg body weight 1,25(OH)_2_-vitamin D_3_ (Sigma D1530) in ethanol 1% in NaCl 0.9%. The dose and application route was the same as previously used in our laboratory. Control mice were injected with 1% (v/v) ethanol in NaCl 0.9%. Mice were sacrificed 6h after injection.

For parathyroid hormone treatment, male C57BL/6N mice aged 12-13 weeks were subcutaneously injected with 80 μg/kg body weight human PTH fragment 1-34 (hPTH1-34) (Sigma P3796) in NaCl 0.9% or NaCl 0.9% alone as vehicle and sacrificed 2h after injection. The dose and application route used was determined from the literature (Kramer et al., 2010).

For estradiol treatment, male C57BL/6N mice aged 16 weeks received 1 daily subcutaneous injection of 15μg 17β-Estradiol (Sigma E8875) in ethanol 0.1% (v/v) in NaCl 0.9% for 5 consecutive days and were sacrificed 4h after the last injection. The dose per body weight and the drug application route was derived from the literature, as shown Abel et al. to induce *Trpv5* expression (Van Abel et al., 2002). Control mice were subcutaneously injected with 0.1% ethanol in NaCl 0.9% for 5 days.

### Mouse dissection

For euthanasia, mice were intraperitoneally injected with 0.1 mg/gBW of ketamine (Ketanarkon 100 Vet., Streuli) and 0.02 mg/gBW of xylazine (Rompun, Bayer), followed by terminal exsanguination by orbital puncture under full anesthesia and /or by cervical dislocation. Organs were collected, kidneys were cut in half, and organs were snap frozen in liquid nitrogen immediately, followed by storage at −80°C until further use.

### RNA extraction

RNA was extracted using TRI reagent (Applied Biosystems by Life Technologies) according to manufacturer’s instructions. RNA pellets were dried and dissolved in RNase-free H_2_O. RNA concentration was measured photometrically using Nanodrop (Nanodrop 2000, Thermo Fisher Scientific, Waltham, MA, USA). RNA A260/A280 ratio was assessed, and each RNA sample was visualized on a 1% agarose gel. RNA was reverse transcribed to cDNA using the PrimeScript RT reaction kit (Takara Bio Inc, Otsu, Japan). RNA input quantities per sample were 1-2 μg for bone, 500ng for kidney or 1μg of MLO-Y4 RNA. The resulting cDNA mix was diluted 2-12x depending on tissue type.

### qPCR

For quantitative gene transcript expression analysis, 2uL of cDNA was used for SYBR Green qPCR (Applied Biosystems by Life Technologies) on a 7500 Fast machine (Applied Biosystems). Samples were run in triplicates in 20uL total volume for each gene, and actin or GAPDH was used for normalization. Melting curves were obtained for every run. Program settings were: 95°C during 20 seconds, 40 cycles (95°C 3 sec, 60°C 30 sec), and for melting curve stage: 95°C 15sec, 60°C 1min, rising at 1% ramp speed to 95°C (15sec), 60°C 15sec. Data were analyzed using the delta-delta CT method. Primers were ordered from Microsynth (Switzerland), and sequences are shown in Table 1. All amplified products were visualized on agarose gels.

**Table 1:**
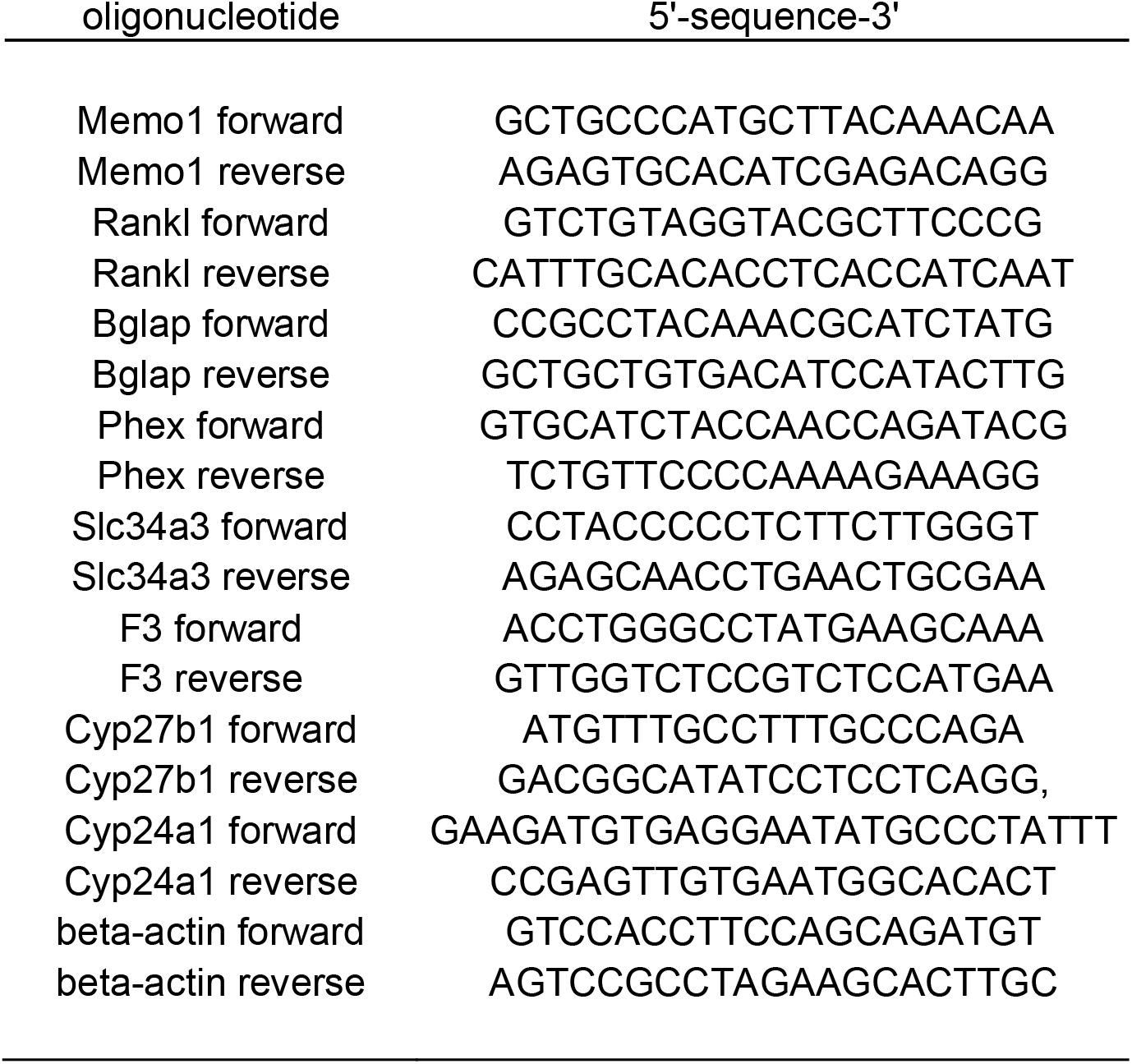
Primers used for qPCR.

### Data analysis

Data from experiments with 2 independent groups were analyzed by t-test or Mann-Whitney U test. For comparison of 3 groups, Kruskal-Wallis test was used with Bonferroni’s Multiple Comparison post-test. All statistical analyses were conducted using GraphPad PRISM 5.03. Two-sided p-values < 0.05 were considered significant.

## Results

*Memo1*-deficient mice resemble by some traits *Klotho* mutant and *FGF23* KO mice (Haenzi et al., 2014), and the promoters of *Klotho* and *FGF23* harbor regulatory sequences that can be bound by vitamin D receptors (Forster et al., 2011, Orfanidou et al., 2012). For these reasons, we determined the regulation of *Memo1* gene expression by minerals and calciotropic hormones. We have previously shown that MLO-Y4 osteocyte-like cell line expressed *Memo1* transcripts and protein (Moor, 2018).

### *Memo1* is diminished by 1,25(OH)_2_-vitamin D_3_ in vitro but not in vivo

We investigated the effects of stimulation with 10nM 1,25(OH)_2_-vitamin D_3_ on gene expression in MLO-Y4 cells (Fig 1). Known vitamin D-dependent transcripts were first assessed. Transcripts of *Cyp24a1* encoding a vitamin D inactivating hydroxlyase (Fig 1a) and of osteoclast regulator *Rankl* (Fig 1c) were increased upon 1,25(OH)_2_-vitamin D_3_ treatment, whereas transcripts of FGF23 regulator *Phex* and expression of bone-derived hormone *osteocalcin* / *bone gamma-carboxyglutamate* (*Bglap*) were diminished (Fig 1b, 1d). *Memo1* transcripts were reduced by 20% upon 1,25(OH)_2_-vitamin D_3_ treatment (Fig 1e).

**Fig. 1.**
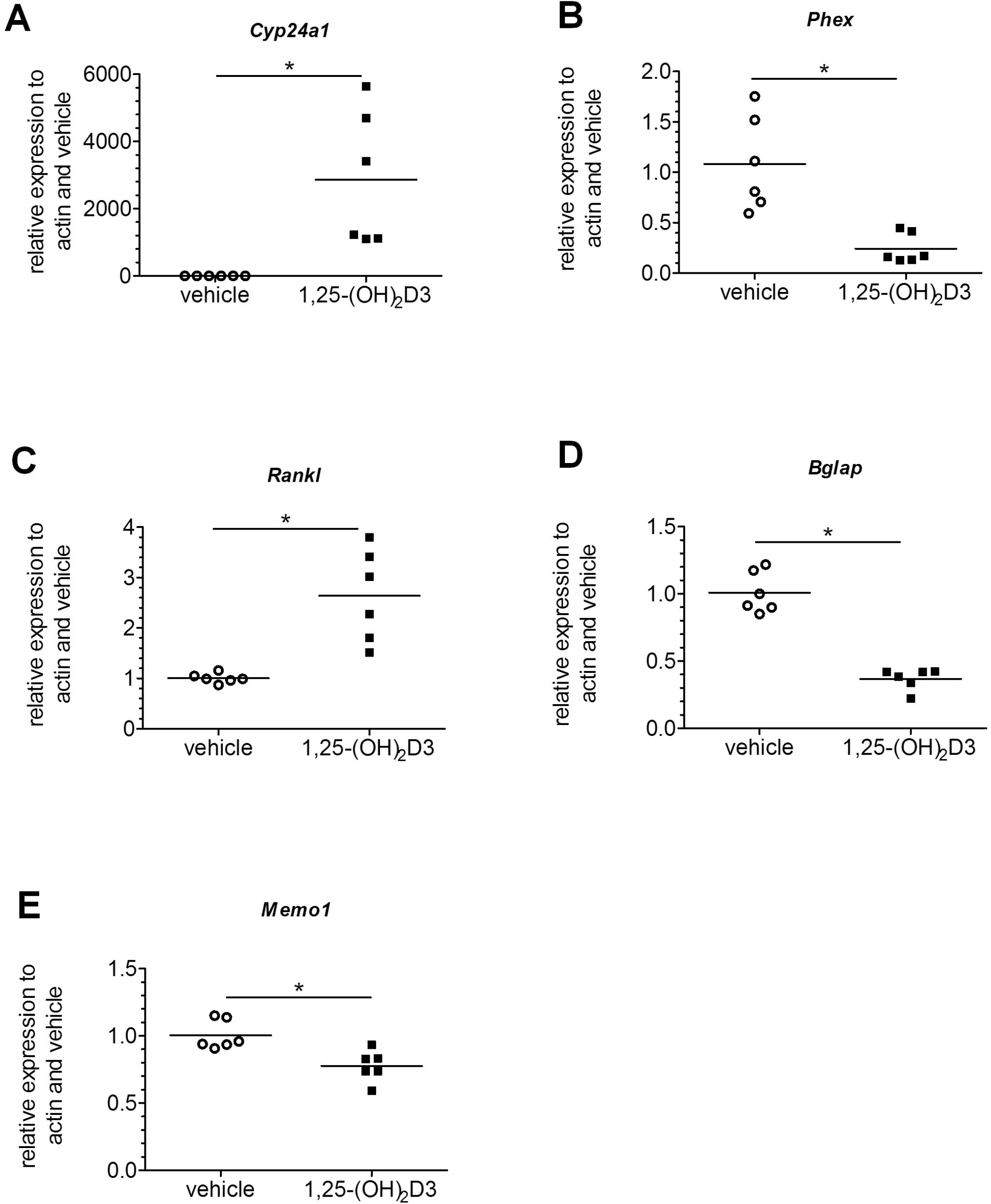
Transcriptional responses to 1,25(OH)_2_-vitamin D_3_ in MLO-Y4 osteocyte-like cells. 1,25(OH)_2_-vitamin D_3_ treatment significantly increased *Cyp24a1* transcripts (A), decreased *Phex* transcripts (B), increased *Rankl* transcripts (C), decreased *Bglap* (osteocalcin) transcripts (D), and decreased *Memo1* (E) transcripts in MLO-Y4 cells. n=6 per condition, * p<0.05 (t-test).

Next, we intraperitoneally injected 1,25(OH)_2_-vitamin D_3_ in mice. Six hours post injection we harvested the kidney and tibia of these animals and investigated the mRNA levels of *Memo1*. *Memo1* RNA levels remained unchanged in the kidney (Fig 2a) and tibia (Fig 2b) compared to control animals injected with vehicle only. As experimental controls, we chose *Cyp24a1* and *Fgf23*. Renal transcripts of *Cyp24a1*, the gene encoding the vitamin D inactivating enzyme cytochrome P450 24a1, were upregulated by 1,25(OH)_2_-vitamin D_3_ compared to vehicle (Fig 2c). In addition, expression of *Fgf23* in the tibia was increased by 1,25(OH)_2_-vitamin D_3_ (Fig 2d).

**Fig. 2.**
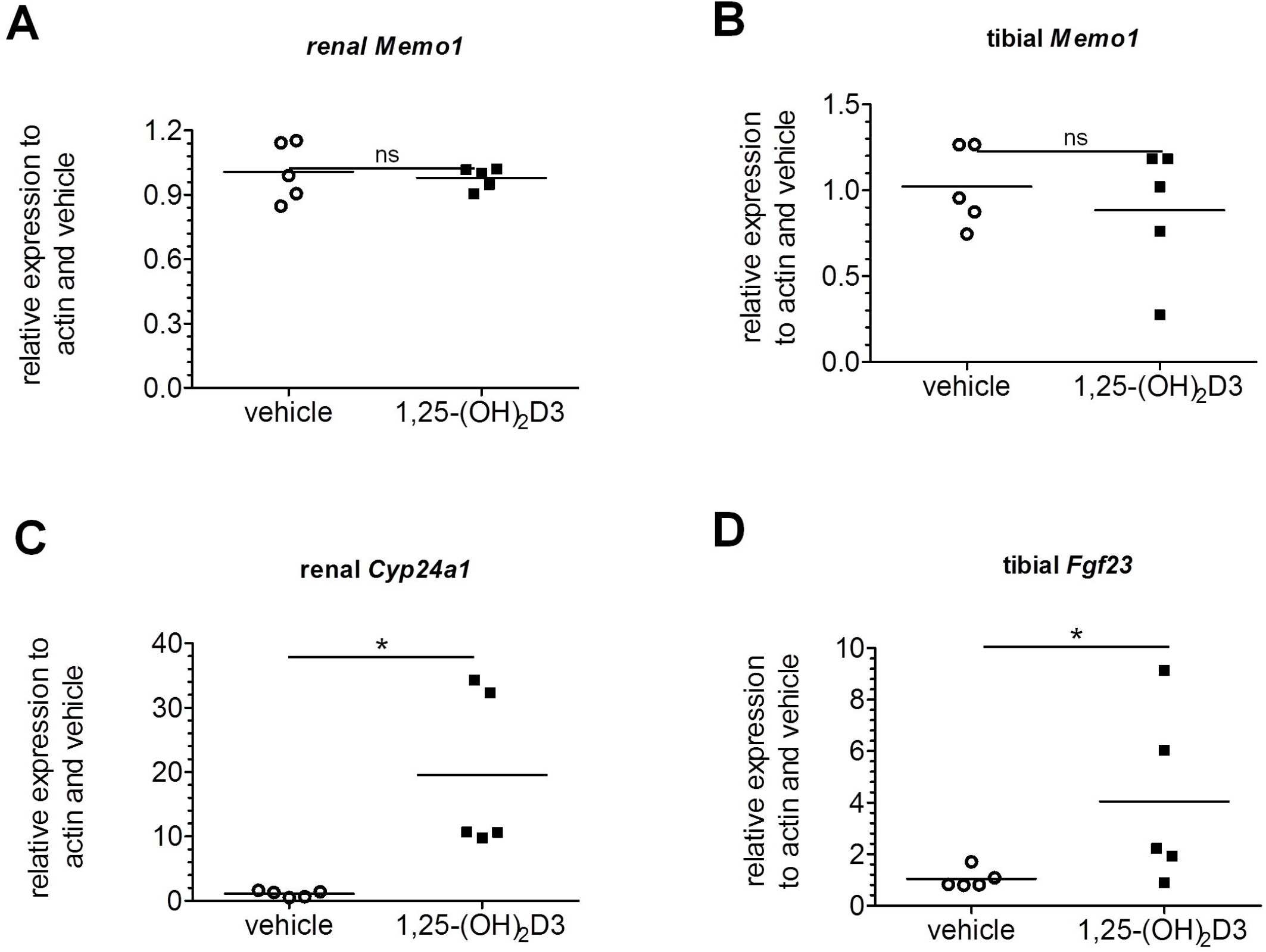
*Memo1* transcripts were not changed by 1,25(OH)_2_-vitamin D_3_ treatment in wildtype mice. *Memo1* transcripts were assessed in kidney (A) and tibia (B) and were indifferent 6h after 1,25(OH)_2_-vitamin D_3_ injection, whereas renal *Cyp24a1* transcripts were over 10-fold increased in comparison to vehicle control (C). *Fgf23* gene expression in tibia was increased by 1,25(OH)_2_-vitamin D_3_ compared to vehicle (D). n=5 per condition, * p<0.05 (Mann-Whitney-U test).

### *Memo1* is not regulated by dietary calcium

Next, we determined the effect of varying dietary calcium content for 7 days on *Memo1* gene expression. RNA was obtained from a previous experiment performed in our lab (366). In the kidney (Fig 3a) and in the tibia (Fig 3b) of these mice exposed to three different calcium-containing diets (0.17%, 0.82% and 1.69%), no change in *Memo1* gene expression was observed. A change in gene expression of *Casr* encoding the calcium-sensing receptor serves as an experimental control for the dietary intervention and was reported for the samples we used in (O’Seaghdha et al., 2013).

**Fig. 3.**
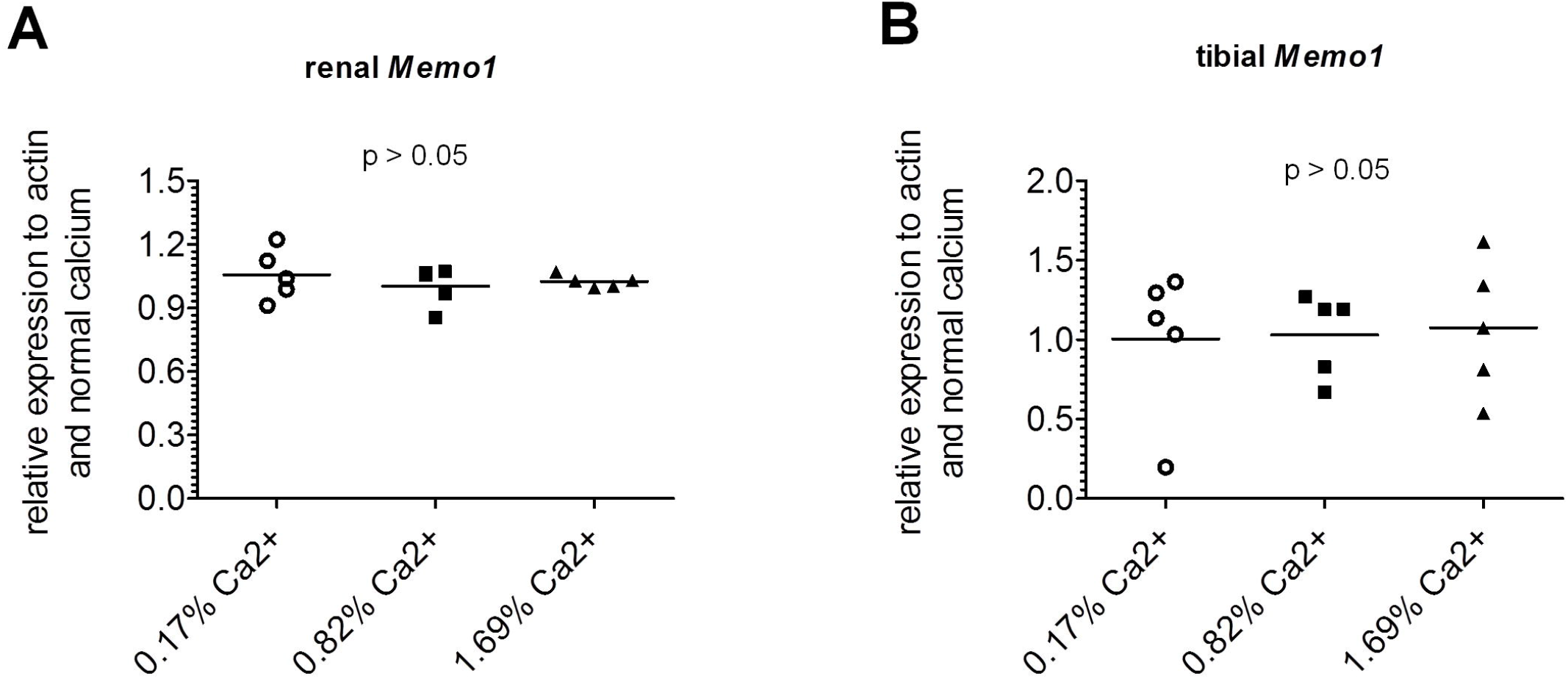
*Memo1* transcript levels were not influenced by varying dietary calcium contents. *Memo1* transcripts in kidney (A) and tibia (B) were not significantly affected by different dietary calcium intake; n=5 per diet (Kruskal-Wallis tests).

### *Memo1* is not regulated by dietary phosphate

We investigated the influence of different systemic phosphate loads on *Memo1* expression. We showed that different dietary phosphate contents (0.2%, 0.8%, 1.5%) did not significantly affect *Memo1* transcript levels in kidney (Fig 4a) or in the tibia (Fig 4b). As an experimental control gene we used renal transcripts of *Slc34a3* encoding sodium-dependent phosphate transporter type 2c (NaPi2c). Renal *Slc34a3* was increased, as expected, under low phosphate and decreased under high phosphate diets (Fig 4c).

**Fig. 4.**
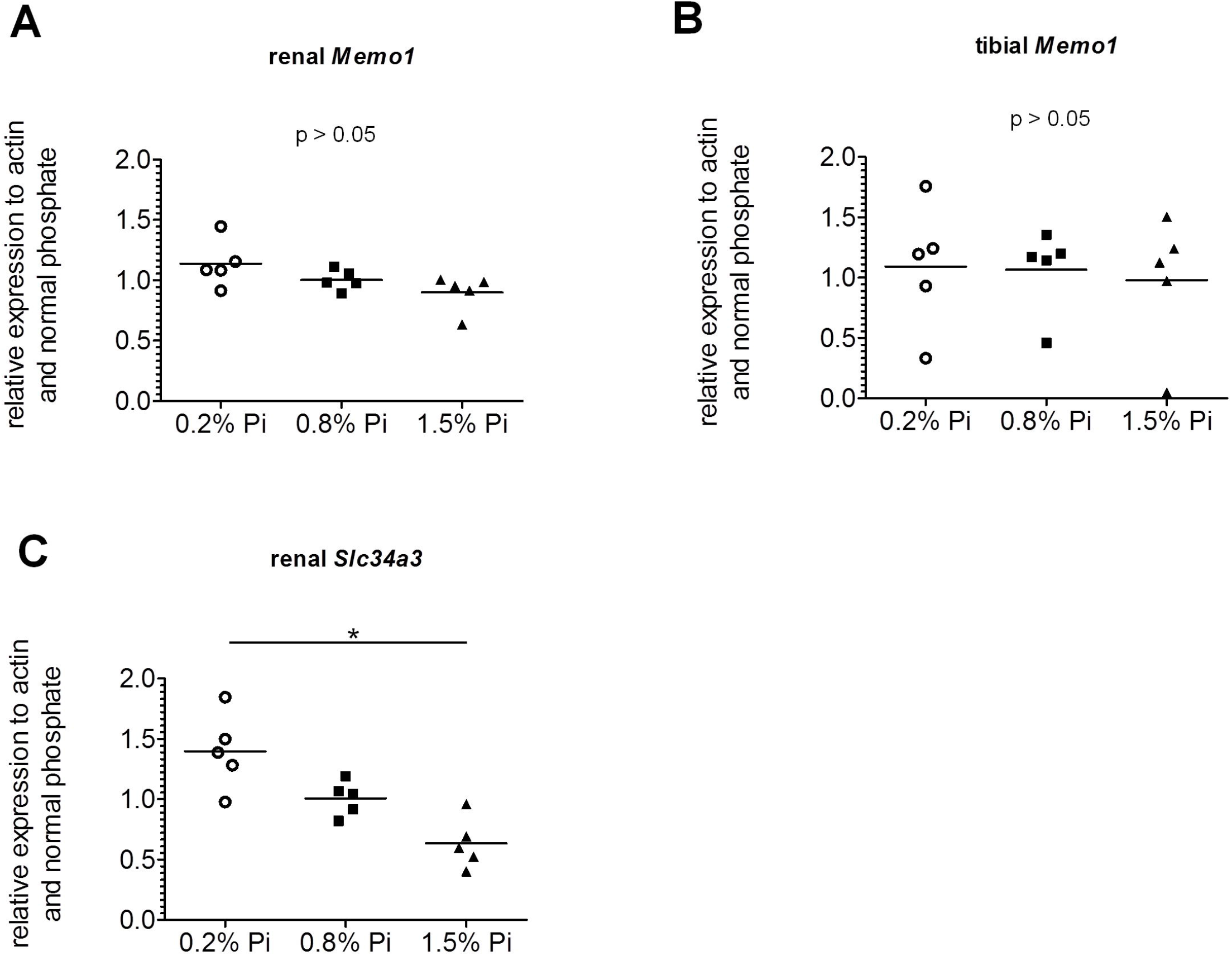
*Memo1* transcript levels were not affected by varying dietary phosphate contents. *Memo1* transcripts abundance in kidney (A) and in tibia (B) were not significantly changed by dietary phosphate contents. Neither bone *Memo1* (B). Renal *Slc34a3* transcripts were used as an experimental control gene and were affected by dietary phosphate content (C). N=5 per condition; *p<0.05 (Kruskal-Wallis test with Dunn’s multiple comparisons correction).

### *Memo1* unchanged by PTH

To determine the effect of PTH on *Memo1*, human PTH fragments 1-34 were subcutaneously injected to wildtype mice, and the animals were euthanized after 2h. *Memo1* gene expression in kidney (Fig 5a) or in tibia (Fig 5b) remained unchanged upon PTH treatment. Transcripts of *Cyp27b1*, the gene coding for the renal vitamin D activating enzyme cytochrome P450 27b1 were increased upon PTH compared to NaCl 0.9%-treated controls (Fig 5c).

**Fig. 5.**
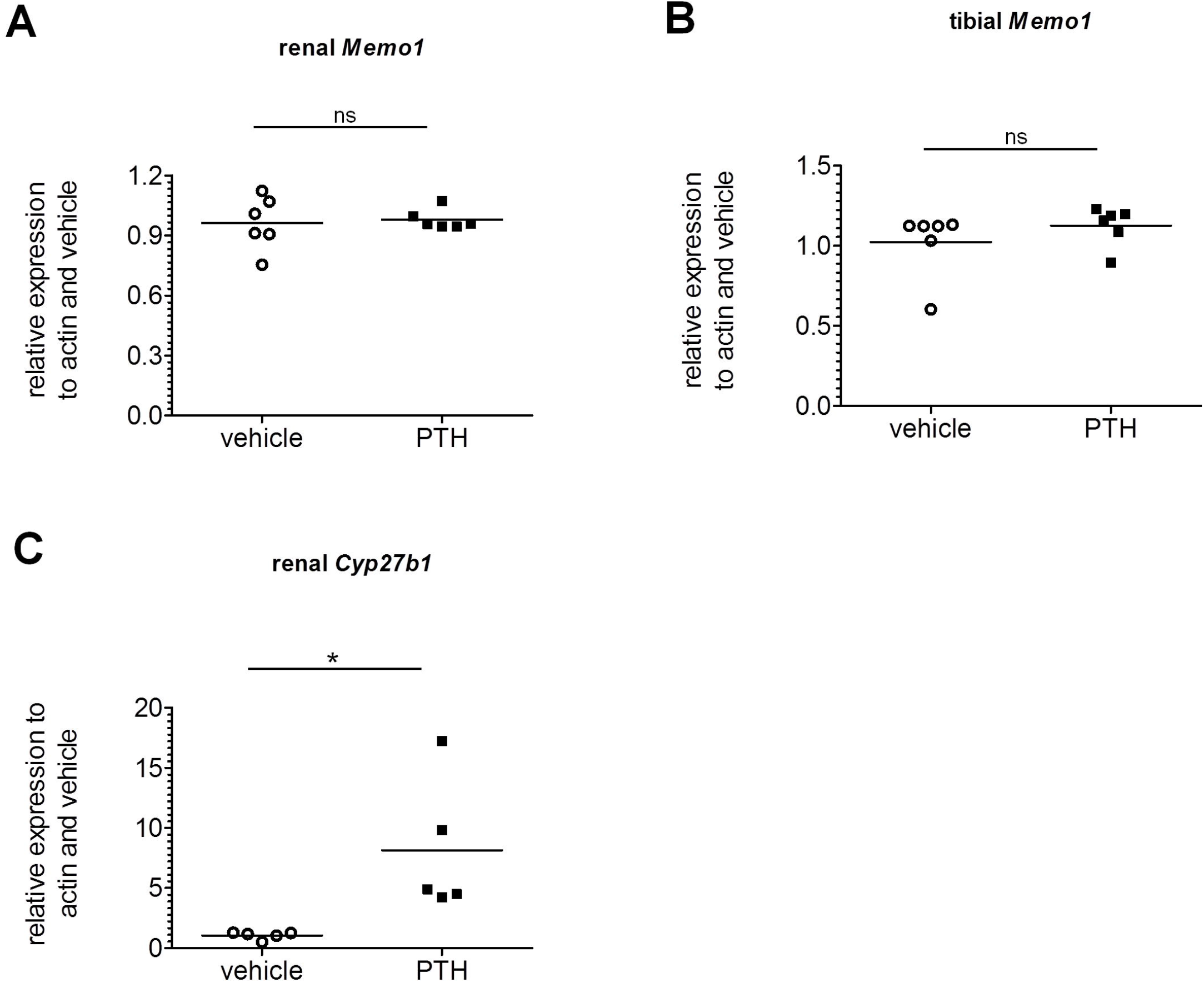
*Memo1* gene expression remained unchanged upon PTH treatment in wildtype mice. Human PTH1-34 or NaCl 0.9% vehicle was injected 2h prior to dissection, and transcripts of *Memo1* in kidney (A) and tibia (B) were unchanged between experimental conditions. Transcripts of *Cyp27b1* (C) were increased by PTH1-34. n= 6 per condition; ns, not significant; * p<0.05 (Mann-Whitney-U test).

### *Memo1* unchanged by estradiol

As sex hormones exert effects on both renal calcium transport proteins (Van Abel et al., 2002) and FGF23 in the bone (Carrillo-Lopez et al., 2009), we tested if *Memo1* is a target gene induced by estradiol. We subcutaneously injected 17β-estradiol once daily over 5 days. This induced the expression of the control gene *F3* encoding coagulation factor III (Fig 6c), but gene expression of *Memo1* in the kidney (Fig 6a) and in the bone (Fig 6b) both remained unchanged compared to mice injected with vehicle.

**Fig. 6.**
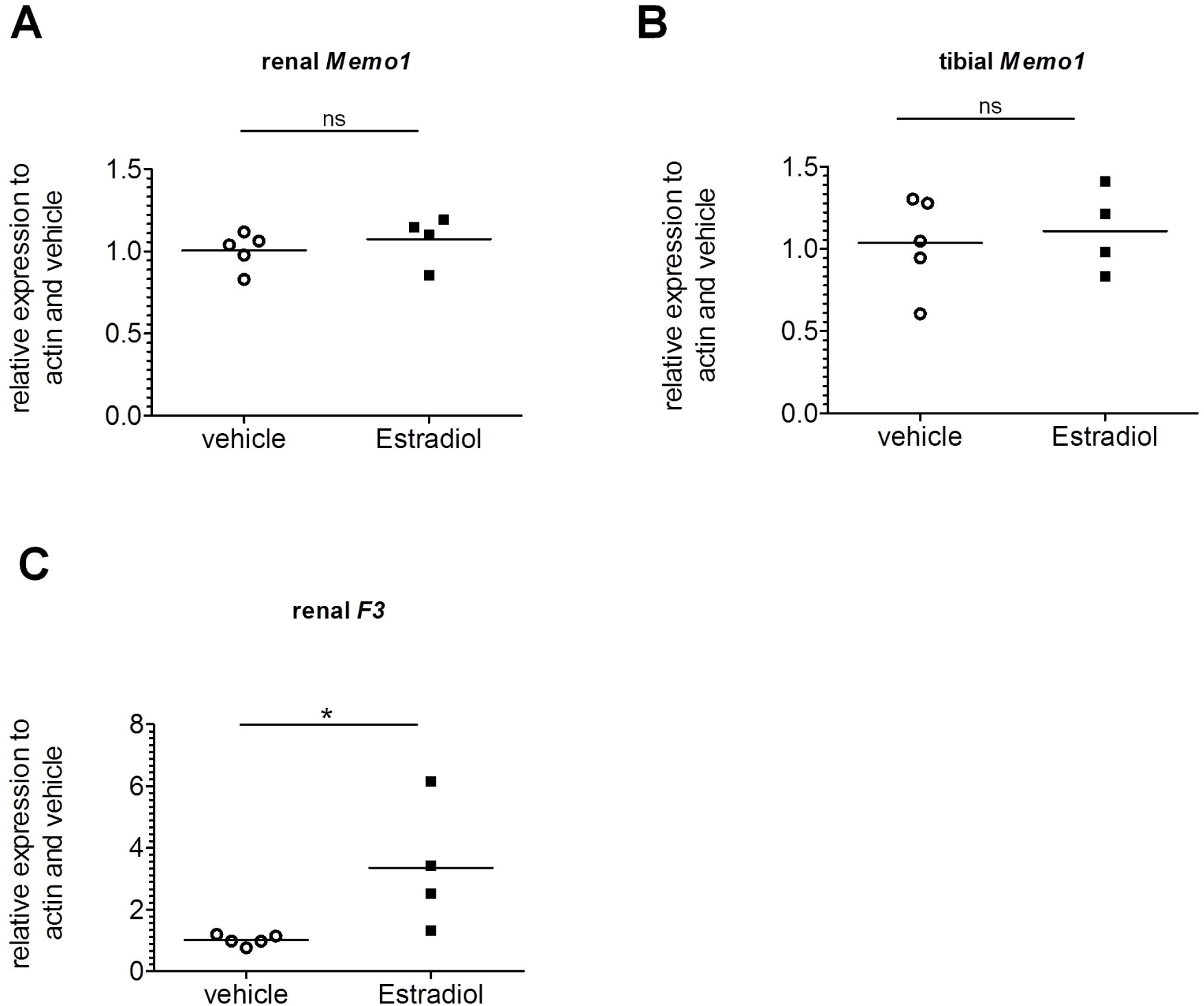
*Memo1* transcripts remained unchanged upon 17β-estradiol treatment. *Memo1* transcripts assessed by qPCR in kidney (A) and tibia (B) were unchanged after 5 daily subcutaneous injections of 17β-estradiol compared to vehicle. Renal gene expression of tissue factor *F3* was increased by 17β-estradiol (C). n=4 estradiol and n=5 control condition. ns, not significant; * p<0.05 (Mann-Whitney-U test).

To summarize, we found a small but significant decrease in *Memo1* expression upon 1,25(OH)_2_-vitamin D_3_ exposure *in vitro*, but we failed to detect any major regulation of *Memo1* transcript abundance upon mineral load or calciotropic hormone treatment *in vivo*.

## Discussion

MEMO1 is expressed in the kidney where it plays an intrarenal role in the regulation of calcium transporters (Moor MB, 2018). In the bone, MEMO1 is expressed in all cell types (Moor, 2018), but its precise bone-specific function remains elusive. Here we tested the hypothesis whether *Memo1* is regulated by key players in mineral homeostasis such as calciotropic hormones or dietary calcium or phosphate. As a readout, we chose *Memo1* gene expression in an osteocyte-like cell line, and in bone and kidney tissues. For each intervention, an experimental control gene was assessed and revealed effects similar as shown before by others.

We observed that *Memo1* gene expression was diminished in the osteocyte-like cells upon 1,25(OH)_2_-vitamin D_3_ treatment. However, in bone and tissues, we failed to detect any effect on *Memo1* by all interventions that we studied. This shows a major difference between *Memo1* and the most studied contributors to mineral homeostasis. As examples in the kidney, Type II sodium-dependent phosphate cotransporters are regulated by dietary phosphate supply (Bourgeois et al., 2013), and gene expression of *Trpv5* encoding a renal calcium transport protein is controlled by 1,25(OH)_2_-vitamin D_3_ (Hoenderop et al., 2001). As examples in the bone, *Fgf23* expression is stimulated by 1,25(OH)_2_-vitamin D_3_ (Liu et al., 2006) or PTH (Kawata et al., 2007), while dietary phosphate restriction or renal phosphate-wasting disorders reduce *Fgf23* expression (Vervloet et al., 2011, Schlingmann et al., 2016, Ansermet et al., 2017). Even intravenous calcium loading in rats increased *Fgf23* expression in bone and hormone concentrations in the serum (Shikida et al., 2018).

This study contains some limitations: The current interventions were confined to analysis of gene expression, but we did not directly assess *Memo1* promoter activity using a reporter construct. Such an approach would more sensitively discriminate and would allow to identify possible response elements in the *Memo1* promoter. However, we argue that a physiologically relevant regulation of *Memo1* gene expression, if present, should have already been visible using the approach undertaken.

Further, we have assessed a single but physiologically reasonable time point, and only a narrow selection of tissues and cells. In addition to bone and kidney, the intestine would be another major turnover place for minerals. *Memo1* expression and potential regulation in healthy intestine has not been investigated so far. In colorectal cancer cells *Memo1* promoter activity is increased in response to the transcription factors Aryl hydrocarbon receptor / Aryl hydrocarbon receptor nuclear-translocator complex, indicating some intestinal disease relevance (Bogoevska et al., 2017).

Finally, as Memo is a redox enzyme with incompletely understood reaction partners and substrates (MacDonald et al., 2014), future studies assessing posttranslational regulation such as by oxidative modification of MEMO1 protein, subcellular localization, or changes in its putative enzymatic activity may be helpful to investigate a regulation of Memo1.

To conclude, besides a minor effect in bone cells stimulated with 1,25(OH)_2_-vitamin D_3_, we did not detect a major regulation of *Memo1* gene expression upon minerals and calciotropic stimuli in bone and kidney, two organs relevant for mineral homeostasis. Further studies inquiring the regulation of this and similar genes may contribute to the understanding of the regulation of mineral homeostasis in health and renal and bone diseases.

## Author participation

OB conceived the project. MBM and OB participated in experimental design. MBM performed experiments. MBM and OB participated in data analysis and interpretation. MBM wrote the manuscript. All authors critically read and commented on the manuscript and agreed to manuscript submission.

## Acknowledgments

OB and MBM’s work was supported by the Swiss National Science Foundation through the special program NCCR Kidney.CH and by unrestricted grants from the Association pour l’Information et la Recherche sur les maladies rénales Génétiques (AIRG)-Suisse and from the Novartis Foundation. The authors are thankful to Candice Stoudmann and Finola Legrand for assisting with an experiment.

